# Key Drivers of Ecological Assembly in the Hindgut of Atlantic Cod (Gadus morhua) when Fed with a Macroalgal Supplemented diet – How Robust Is the Gut to Taxonomic Perturbation?

**DOI:** 10.1101/2021.08.24.457479

**Authors:** C. Keating, M. Bolton-Warberg, J. Hinchcliffe, R. Davies, S. Whelan, A. H. L. Wan, R. D. Fitzgerald, S. J. Davies, C. J. Smith, U. Z. Ijaz

**Affiliations:** Division of Microbiology, School of Natural Sciences, National University of Ireland Galway, Ireland, H91 TK33; Water and Environment Group, Infrastructure and Environment Division, James Watt School of Engineering, University of Glasgow, Glasgow, United Kingdom, G12 8LT; Institute of Biodiversity, Animal Health & Comparative Medicine, University of Glasgow, Glasgow G12 8QQ, UK; Carna Research Station, Ryan Institute, National University of Ireland Galway, Carna, Co. Galway, Ireland, H91 V8Y1; Department of Biological and Environmental Sciences, University of Gothenburg, Gothenburg, Sweden; AquaBioTech Group, Central Complex, Naggar Street, Targa Gap, Mosta, MST 1761, Malta G.C.; Irish Seaweed Research Group, Ryan Institute and School of Natural Sciences, National University of Ireland Galway, Ireland, H91 TK33; Aquaculture and Nutrition Research Unit, Carna Research Station, Ryan Institute and School of Natural Sciences, National University of Ireland Galway, Carna, Co. Galway, Ireland, H91 V8Y1; Department of Animal Production, Welfare and Veterinary Science, Harper Adams University, Newport, Shropshire, UK, TF10 8NB

## Abstract

The microbiota in the gastrointestinal tract of any species is shaped by internal and external cues in addition to random events which can be difficult to disentangle from a range of interacting variables. Estimating ecological assembly processes can help elucidate these factors. In our study, farmed Atlantic cod (*Gadus morhua*) were fed a diet of 10% macroalgae supplement (*Ulva rigida* species [ULVA] or *Ascophyllum nodosum* [ASCO] or a non-amended control diet [CTRL]) over a 12-week period and the ecological assembly processes quantified. The CTRL samples showed selection (variable selection - divergence in communities related to selective environmental conditions) as a key assembly process, while dispersal limitation (limited movement results in divergent communities through stochastic processes) was a driver of the gut microbiome for fish fed the macroalgae supplemented diet at Week 12 (i.e., ASCO and ULVA). Fish from the ASCO grouping diverged into ASCO_N (normal) and ASCO_LG (lower growth), where ASCO_LG individuals found the diet unpalatable. The recruitment of new taxa over time was altered in the ASCO_LG fish, with the gut microbiome showing phylogenetic under dispersion (nepotistic recruitment of species). Finally, the gut microbiome (CTRL and ULVA only) showed increasing robustness to taxonomic disturbance over time and an increase in functional redundancy. This study advances our understanding of the ecological assembly and succession in the hindgut of juvenile Atlantic cod across dietary regimes. Understanding the processes driving ecological assembly in the gut microbiome, in fish research specifically, could allow us to manipulate the microbiome for improved health or resilience to disease for improved aquaculture production.

## 1. Introduction

Aquaculture has become the fastest-growing food sector this century, surpassing over 82 million tonnes in seafood production in 2018^1^ with its contribution to global seafood production exceeding 45%^1^. At present, however, Atlantic cod (*Gadus morhua*) is primarily harvested through capture fisheries owing to species-specific production bottlenecks in aquaculture, leading to reduced profitability^2,3^. Such bottlenecks include poor larval survival rates^4^, early sexual maturation (reduced fillet yield)^5^, and reduced fish growth^6^. A review of these challenges is noted here^7^. An avenue that is being increasingly explored is the addition of aquafeed ingredients that provide additional health benefits to the growing fish from a variety of natural sources (e.g., macroalgae supplements^8–10^, immunostimulants^11,12^, pre- and probiotics^13,14^). In conjunction with such diet supplements, it is apparent that the impact of these feeds on the gut microbiome and fish health should be considered.

The fish gut microbiome is increasingly being investigated to elucidate fish condition^15^, response to environmental conditions^16^ and changing diet^17,18^ and has been extensively reviewed elsewhere^15,19,20^. The cod gut microbiome in wild populations has been shown to have limited microbial diversity and a high prevalence of closely related species of microflora, such as *Photobacterium*, in wild populations^21,22^. In our previous work, we have shown that there was high diversity in the gut microbiome of farmed juvenile cod monitored over a 12 week period^23^, and while we also reported a high incidence of *Photobacterium* in the hindgut we observed a temporal decrease in *Photobacterium* relative abundance without indications of any environmental cause. It is unclear if this shift was a result of natural community succession in the developing hindgut microbiome of juvenile farmed fish or some other factor. Little is known about the gut microbiome development in cod, with the limited number of studies having only reported on the larval phase^24^ or wild adults^21^. Moreover, the microbial community assembly mechanisms themselves remain underexplored. Exploration of these mechanisms may offer a realistic opportunity to manipulate the gut microbiome to improve fish condition, increase resilience to disease and mitigate stress under aquaculture conditions.

Ecological theory has been used to describe the mechanisms that shape community assembly, and more typically applied to macroecology (e.g. fish species assembly^25^, specific populations^26^), but is increasingly being used in microbial ecology^27–29^ to determine underlying rules driving the assembly and dynamics of complex microbial communities which is particularly challenging^30^. The gut microbiome is a complex and dynamic ecosystem subject to fluctuating abiotic and biotic conditions. Authors have noted that founder (priority) effects can play a key role in the community assembly and succession, i.e., the species that first colonise will alter the environment and determine the resulting community structure^31,32^. The colonisation of that first species may be random like a lottery^31^, or determined via resource availability and species traits ^33,34^. Assembly processes are broadly described as either random (stochastic) or non-random (deterministic)^30^. Researchers have noted that four assembly processes can occur in microbial communities: neutral and stochastic processes (e.g. probabilistic events such as births, deaths, mutations and ecological drift^35^) or non-probable events driven from niche or deterministic forces (e.g., environmental conditions, species interactions and traits). Neutral theory differs from niche-based theory in the assumption that all species are equal in terms of functional traits, demographic rates and the environment does not select.

To determine the underlying mechanisms of microbial community assembly a range of tools have been described. Many of these rely on ‘null models’. A null modelling approach considers randomising original community structure and then through a statistical framework compares properties of microbiome between the original and randomised communities to elucidate a particular ecological phenomenon. The randomised community is generated in such a way that it mimics a community without the force of a specified assembly process^36^. Deviations from the null model can then be used to predict the processes occurring in the real community. In our previous work, over a twelve-week period, we fed Atlantic cod juveniles (*G. morhua*) a diet of 10% macroalgal supplement (either *Ascophyllum nodosum* [ASCO] or *Ulva rigida* [ULVA] species) or a control non-amended diet^23^. We showed that temporal pressures outweighed the response to diet supplementation, with the gut microbiome of all fish consuming the different diets converging. A subset of fish found one diet unpalatable and showed reduced growth (ASCO_LG) and did not follow the same trend. The ecological drivers of this convergence of the microbiome with time was not understood therefore, in the present study, the aims are to implement a suite of ‘null-modelling’ tools to understand the microbial ecological assembly mechanisms in the gut of Atlantic cod (*G. morhua*) in an experimental feeding trial over a twelve-week period. Specifically, we aim to determine: i) whether the microbial community was driven by stochastic, deterministic forces, niche, or neutral effects, ii) the influence of specific ecological processes on microbial diversity, iii) colonisation and community succession mechanisms and finally, iv) the resilience of the gut microbial function to taxonomic perturbation. We hypothesised that the gut microbiome would demonstrate deterministic and niche-based assembly linked to host development, that this would outweigh environmental variables such as feed type, and that individuals with poorer growth (ASCO_LG) would demonstrate vulnerabilities to taxonomic perturbation. This work has significant importance with respect to fish gut microbiome research. More broadly, this work is relevant to areas whereby manipulating the microbial community through the application of novel feed additives and functional supplements is desired.

## 2. Results

It was evident that there were strong temporal pressures on the hindgut microbial community of juvenile Atlantic cod (*G. morhua*), with a change in the microbial community composition over the course of the 12-week trial. The OTUs that best correlated with the temporal community dissimilarities (Bray-Curtis dissimilarity matrix) were *Proteobacteria, Bacteroidetes* and *Firmicutes* species. *Proteobacteria* spp. (OTU_3130 + OTU_307 - *Photobacterium* spp. and OTU_2746 - *Vibrio* sp.) were associated with Week 0 to Week 8 (Figure 1A). While *Bacteroidetes* spp. (OTU_9 + OTU_3586 - *Bacteroides* spp. and OTU_41 + OTU_37 - *Rikenella* spp.) and *Firmicutes* spp. (OTU_1855 + OTU_174 + OTU_33 - *Ruminococcaceae* spp., OTU_2624 - *Lachnoclostridium* sp. and OTU_23 - *Tyzzerella* sp.) correlated with Week 12 samples (Figure 1). The OTU subset that most explained the dissimilarity pattern included all the aforementioned OTUs with the exception of OTU_37 – *Rikenella* sp. (R = 0.902). Notably, a temporal shift in OTUs was not observed in the ASCO_LG individuals. Predictive functional analysis using PICRUSt2 revealed that some of the detected microbial metabolism pathways had significantly changed (greater than Log2 fold) from Week 0 to Week 12 (Supplementary Figure 1). This included the detection of pathways related to lysine biosynthesis and degradation and methane metabolism at Week 12, which were not present at Week 0.

**Figure 1.**
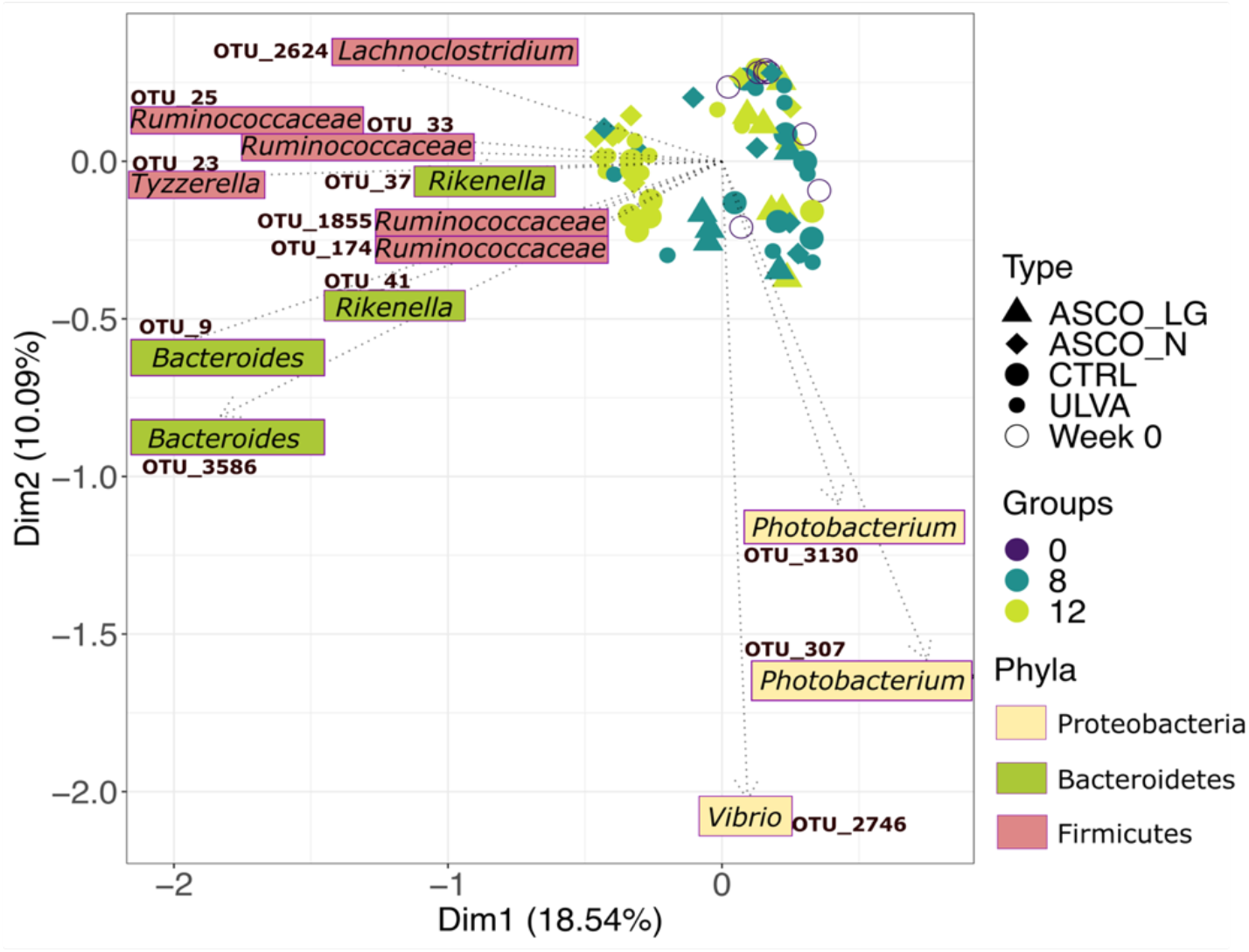
Principal coordinate analysis (PCoA) using Bray-Curtis dissimilarity of the gut microbiota of juvenile Atlantic cod (*Gadus morhua*) community composition based on Time in weeks (*Groups*: 0, 8, 12) and the dietary treatments (Week 0, CTRL, ULVA, ASCO_N and ASCO_LG). The ordination is constrained by super-imposing the subset of OTUs which roughly explained the same beta-diversity between samples as the full dataset of OTUs. Deviance in ordination space is explained by 24.5%. Variation accounted for by *Type*; R^2^ = 0.098, *** P = 0.001. Variation accounted for by *Groups*; R^2^ = 0.08644, *** P = 0.001.

### 2.1. Characterisation of the microbial community assembly mechanisms in the hindgut of juvenile Atlantic cod (Gadus morhua)

Dissimilarity was observed between Week 0 and Week 8 treatments (CTRL, ULVA, ASCO_N and ASCO_LG) in pair-wise comparisons with ^*q*^*d* values > 0.65 (Figure 2Ai). This trend in temporal dissimilarity was also observed between the Week 8 and Week 12 groups (CTRL, ULVA, ASCO_N, and ASCO_LG). However, the ASCO_N and ASCO_LG groups showed less dissimilarity over time with ^*q*^*d* values < 0.5 between Week 8 and Week 12, as compared to the other temporal comparisons. For all groups at q values close to 0 (ignoring relative abundances), dissimilarity was close to the null expectation (explained by random assembly). As q values increased (reflecting the complexity of OTU abundance) dissimilarity was further from the null expectation (explained by deterministic forces). The hindgut microbial community showed a trend towards increasing neutrality (Figure 2B). Week 12 treatment groups were significantly more neutral than Week 0 (P<0.05). However, ASCO_LG samples showed an increase towards niche assembly processes over time, though values remained lower than Week 0.

**Figure 2.**
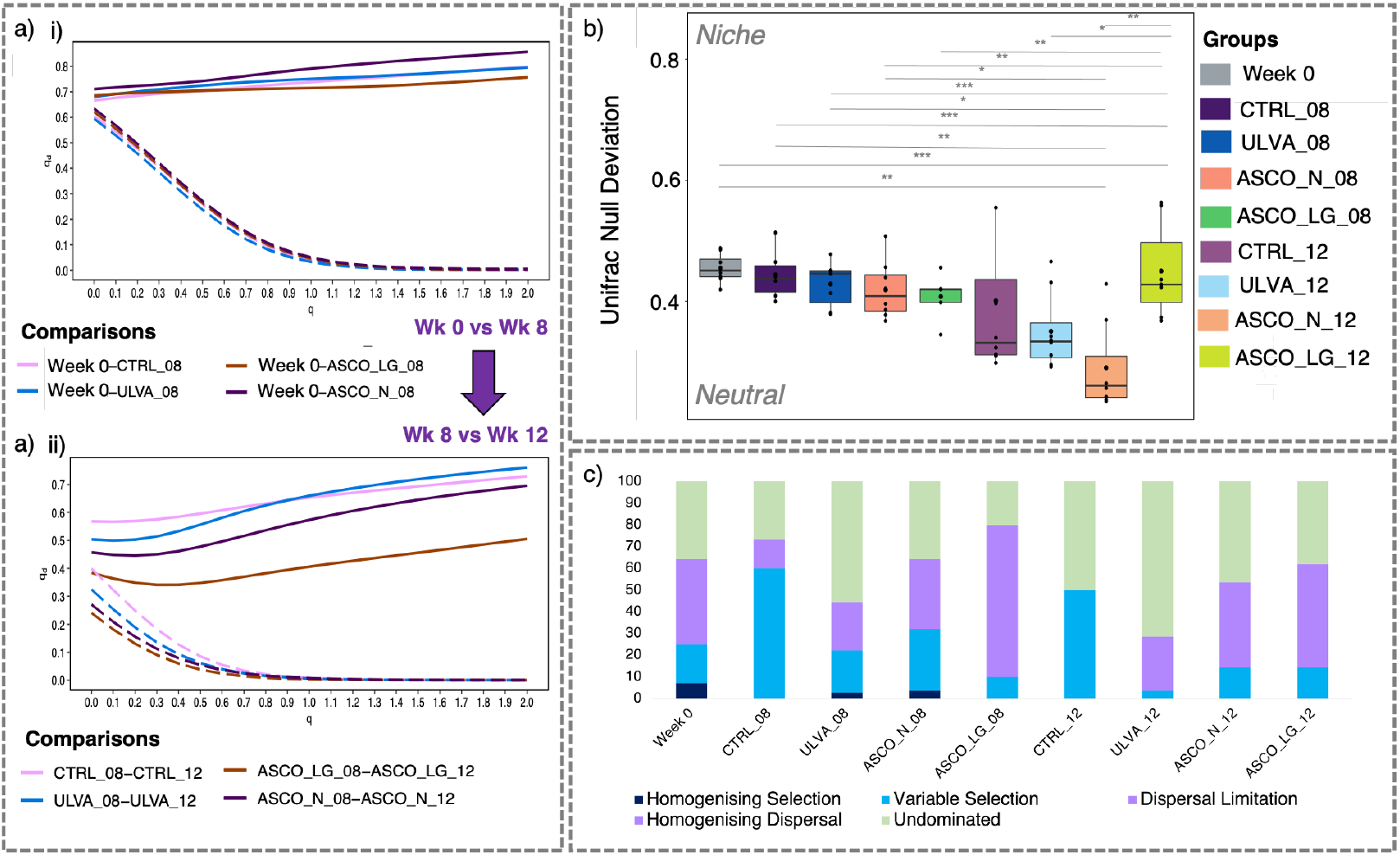
Collated figures exploring the influence of stochastic, deterministic, niche and neutral ecological assembly and quantitative measures of specific ecological processes occurring in the hindgut microbial community of juvenile Atlantic cod (*Gadus morhua*). **a)** *Observed* Hill-based dissimilarity -qd (solid lines) and the null expectation (dashed line) based on 999 randomisations for treatment groups’ pair-wise comparisons; **(i)** Week 0 versus Week 8 treatment groups (CRTL, ULVA, ASCO_N and ASCO_LG); **(ii)** Week 8 versus Week 12 treatment groups (CTRL, ULVA, ASCO_N and ASCO_LG. The x-axis can be interpreted as follows q = 0; presence/absence i.e. Jaccard index, q = 1; OTUs abundances are weighted i.e., Bray-Curtis and q = >1; OTUs with greater relative abundance have increased weighting; **b)** The relative changes in niche and neutral processes assessed using deviations from phylogeny abundance-weighted – Generalised Unifrac beta null model for treatment groups (CTRL, ULVA, ASCO_N and ASCO_LG) over time (Week 8 and Week 12) and including the base-line time-point Week 0. Lines connecting categories shows were significant (ANOVA) with * (p < 0.05), ** (p < 0.01), or *** (p < 0.001). **c)** Stacked bar-chart showing the percentage contribution of homogenising selection, variable selection, dispersal limitation, homogenising dispersal and undominated microbial community assembly processes for Week 0 and the treatment groups (CTRL, ULVA, ASCO_N and ASCO_LG) over time (Week 8 and Week 12). Note that the y-axes values in each plot in the collated figure differ.

At Week 0, dispersal limitation (limited dispersal or historical contingency), variable selection (diverging communities), and homogenising selection (converging communities) accounted for 39, 18, and 7%, respectively, of the assembly processes (Figure 2C). Undominated assembly (i.e., no ecological process dominated) accounted for 36% at Week 0. In the control group [CTRL], variable selection increased to 60% at Week 8. At Week 12, variable selection and undominated both equated to 50% contribution to community assembly in the CTRL treatments. In contrast, the macroalgal diet treatments showed reduced variable selection (4-14%) at Week 12 and increased contribution of dispersal limitation assembly process (25-48%). This was particularly apparent in the ASCO_LG fish where dispersal limitation accounted for 70% of community assembly at Week 8 and 48% at Week 12. Speciation and homogenising selection were not observed as important assembly processes in the cod hindgut microbiome.

### 2.2. Examination of priority effects (competitive lottery model) and species succession (phylogenetic recruitment) in the hindgut of juvenile Atlantic cod (Gadus morhua) over time

All clades (groups of taxa) indicated in Figure 2A had OTUs which were “lottery winners” (> 90% abundance of that clade). According to the threshold Verster and Borenstein (2018)^31^ four clades displayed strong “lottery-like” assembly behaviour (winner prevalence > 0.75 and winner diversity > 0.25) indicating winners observed in nearly all samples. These clades were *Alistipes, Cetobacterium, Fusobacterium*, and *Tyzzerella* genera (Figure 3A; Supplementary Table S1). This behaviour changed over time with both winner prevalence and winner diversity decreasing over time. For example, the *Alistipes* clade with high prevalence (>0.75) at Week 0 and diversity (>0.5), decreased from three OTUs (OTU_34, OTU_20 and OTU_3429) to a sole OTU at Week 8 (OTU_3429) and new sole OTU at Week 12 (OTU_62). The *Bacteroides* clade showed near lottery-like status at Week 0 with a high prevalence (>0.75) and diversity of 0.2 at Week 0, winner diversity decreased from two OTUs (OTU_9 and OTU_2969) to one OTU at Week 12 (OTU_9). In contrast, *Rikenella* and *Lachnoclostridium* clades (with >50 prevalence) showed low diversity initially with a single OTU winning (i.e., OTU_41 *Rikenella* sp. and OTU_2624 *Lachnoclostridium* sp.). Diversity of winning OTUs increased over time with the addition of an OTU per clade (i.e., OTU_37 *Rikenella* sp. and OTU_130 *Lachnoclostridium* sp.) at Week 12. Many of the remaining clades showed low diversity with a single OTU indicated as a ‘winner’. Notably, no *Proteobacteria* genera (which includes Photobacterium) conformed to the competitive lottery schema (data not shown).

**Figure 3.**
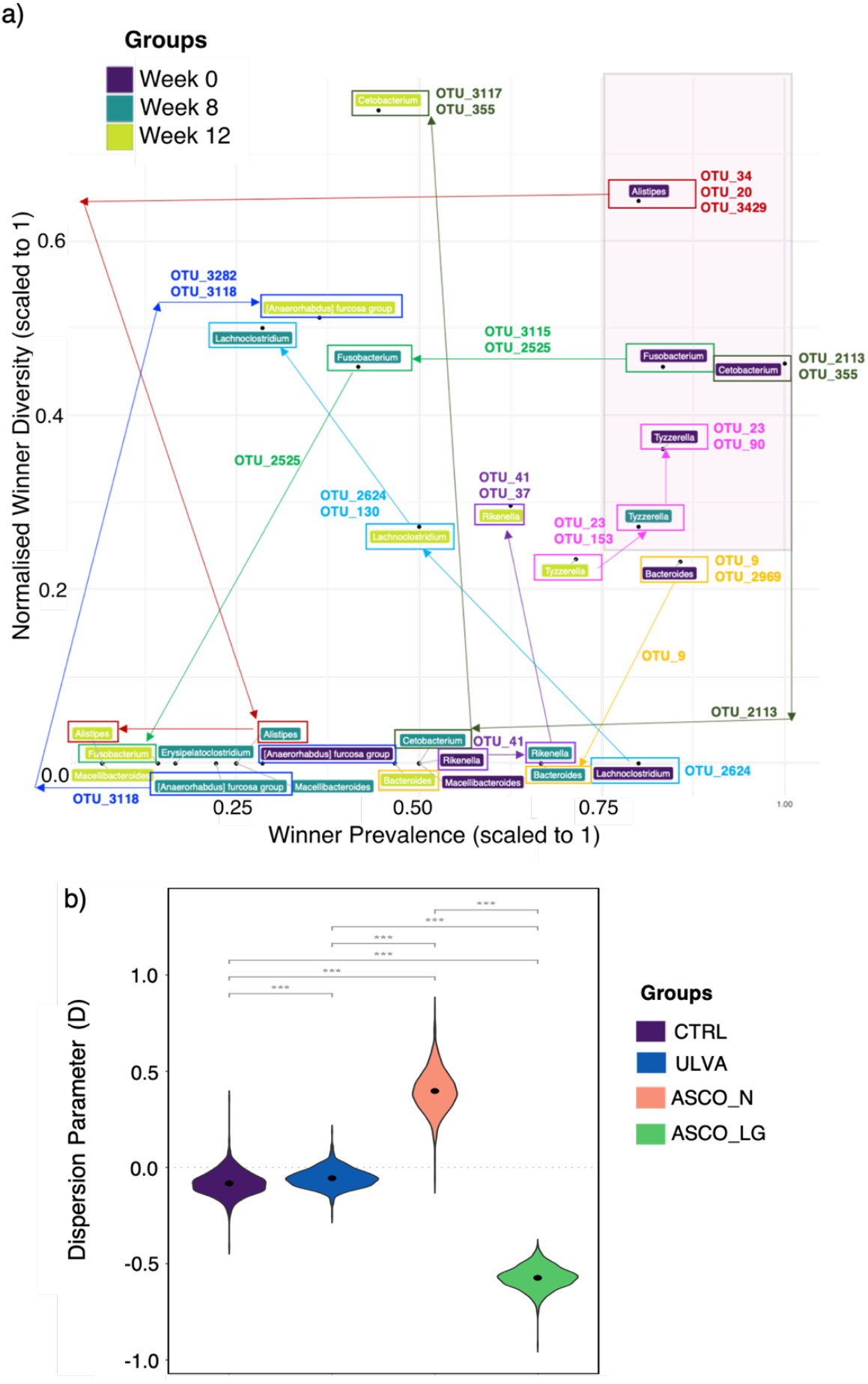
Examination of species colonisation (Competitive lottery model) and species succession (Phylogenetic recruitment model). **a)** Competitive lottery model highlighting the genera with operational taxonomic units (OTUs) showing lottery-like behaviour in Week 0, Week 8, and Week 12 groups. Winner prevalence (percentage abundance of lottery winners across samples) is plotted on the x-axis and winner diversity (count of winning OTUs within a genus across samples) on the y-axis. In the case of high winner diversity, multiple OTUs can show up as winners within a genus, when winner diversity is low a single OTU is highlighted as a winner in the genus. The arrows show the temporal changes in diversity/prevalence of a specific genus. Table of winning OTUs is included in Supplementary File 1. **b)** Phylogenetic recruitment of species over time (Week 8 and Week 12) in the CTRL, ULVA, ASCO_N and ASCO_LG groups. Each violin plot shows the distribution of Dispersion estimates (D) given by logistic error model bootstraps.

The hindgut microbial communities from the CTRL and ULVA dietary fish groups were primarily neutral with respect to phylogenetic dispersion (D=0; Figure 3B), although, there was a high variation in the CTRL D values (ranging from -0.45-0.45). In contrast, the ASCO_N fish demonstrated phylogenetic overdispersion (D>0; D=∼0.45), while the ASCO_LG group were phylogenetically under dispersed (D<0; D=∼-0.45). There was a significant difference between all treatment groups.

### 2.3. Stability of Hindgut Microbiome to Taxonomic Perturbation

In general, attenuation values increased over time from 2.2 at Week 0 to 2.7 at Week 12 (*high values indicate robustness due to functional overlap*). However, these were not found to be statistically significant, and were highest in the ASCO treatments at Week 12 (Figure 4A). Buffering values were in the range of 1.9-2.05 and no temporal or treatment effects were noted (Figure 4B) – *high values indicate functions are balanced across the communities*. We then implemented a principal coordinate analysis (PCOA) using five gene distribution features (GDF). The analysis showed that at Week 0 the microbial community was clustered towards increased unique function abundance and higher average functional redundancy (Figure 4C). The ASCO_LG (Week 8 and Week 12) samples clustered towards increased average genome size, genome size variability, and average functional dissimilarity.

**Figure 4.**
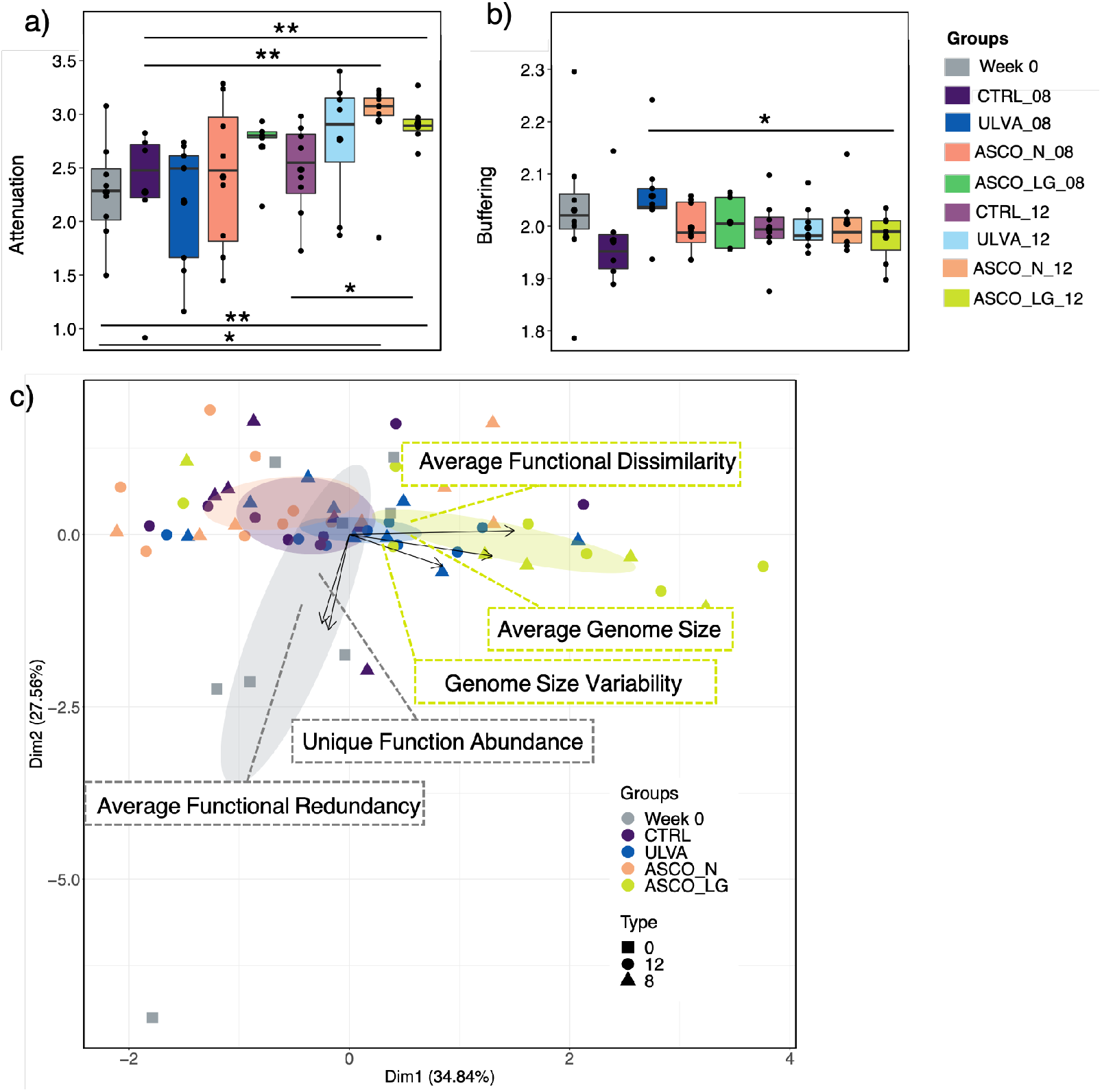
Taxa-function robustness in the Atlantic cod hindgut microbiome compared across Treatments and Time (Groups: Week 0, CTRL_08, CTRL_12, ULVA_08, ULVA_12, ASCO_N_08, ASCO_N_12, ASCO_LG_08 and ASCO_LG_12). **a)** The attenuation values for each group. **b)** The buffering values for each group. **c)** Principal co-ordinate analysis plot of the five gene distribution factors compared across group (Week 0, CTRL, ULVA, ASCO_N and ASCO_LG) over time (Week 0, Week 8 and Week 12). Lines connecting categories shows were significant (ANOVA) with * (p < 0.05), ** (p < 0.01), or *** (p < 0.001).

In general, the attenuation values of specific functions were significantly increased from Week 0 to Week 12 (Figure 5A). In the Supplementary Figure 2, we show the comparisons of the dietary treatments over time with the exclusion of the ASCO group samples. The attenuation values of carbohydrate metabolism increased from ∼1.0 at Week 0 to 2.2 at Week 12 when comparing the over time without the inclusion of any ASCO samples (Supplementary Figure 2). Notably, when making comparisons across all dietary treatments, the attenuation values for functional superpathways related to infectious diseases, cell growth and death, replication and repair increased over time, in all but the ASCO_LG samples (Figure 5A). The dietary subgroup ASCO_LG had increased values for functional superpathways related to the excretory system, translation, transport, and catabolism (Figure 5A). The buffering values for superpathway functions related to metabolism (metabolism of cofactors and vitamins, biosynthesis of other secondary metabolites) decreased over time and across all treatment groups, except ASCO_LG (Figure 5B). A similar trend was also observed in the specific buffering functions related to cell regulation processes (folding, sorting and degradation, replication and repair and signal transduction).

**Figure 5.**
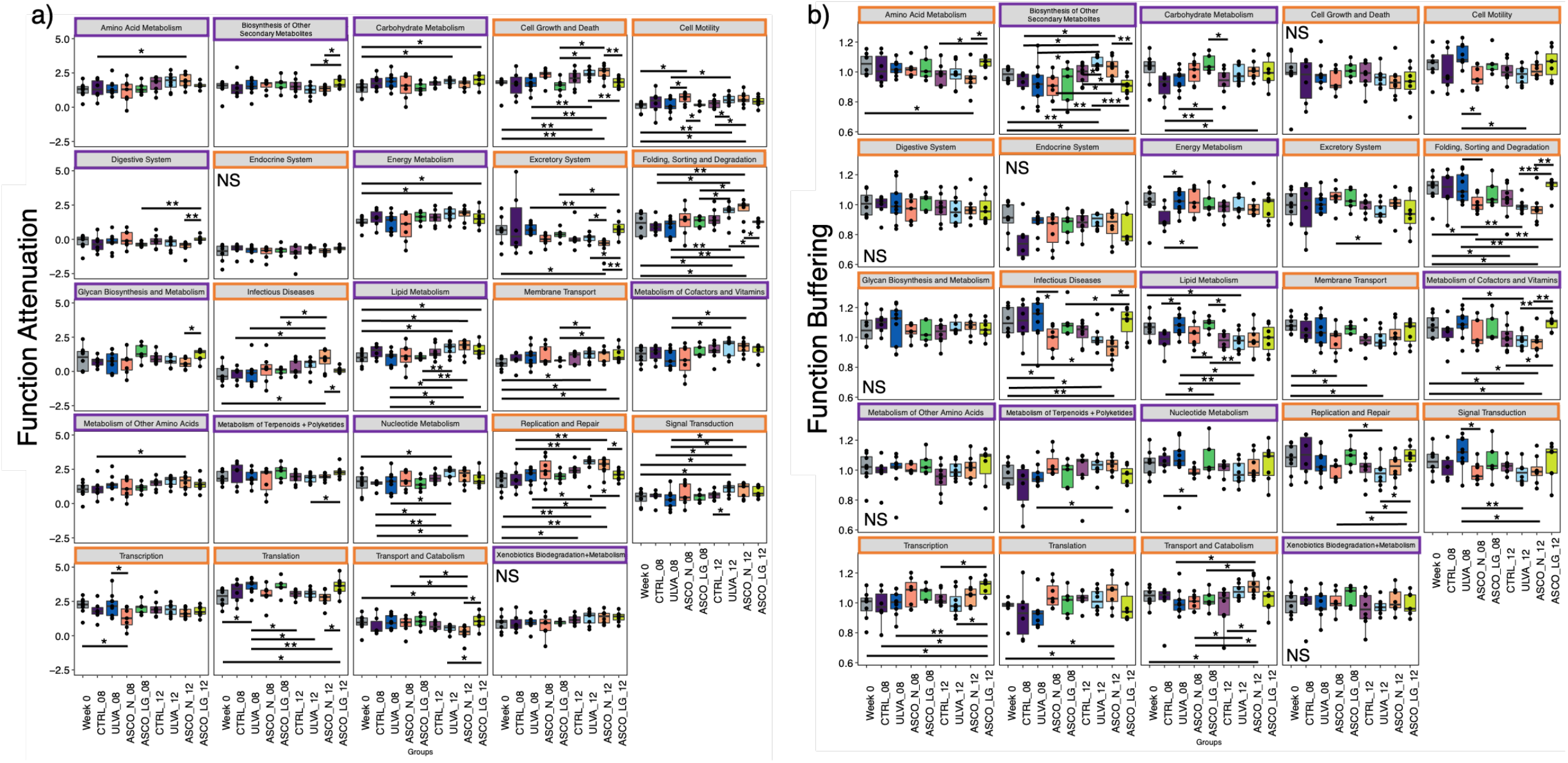
Taxa-function robustness in the Atlantic cod hindgut microbiome compared across treatments and time (Groups: Week 0, CTRL_08, CTRL_12, ULVA_08, ULVA_12, ASCO_N_08, ASCO_N_12, ASCO_LG_08 and ASCO_LG_12) for specific microbial superpathway functions. Functions are highlighted as being involved in metabolism (purple box) or cell functioning (orange box). **a)** The specific-robustness attenuation functions that were statistically significant between groups. **b)** The specific-robustness buffering functions that were statistically significant between groups. Lines connecting categories shows were significant (ANOVA) with * (p < 0.05), ** (p < 0.01), or *** (p < 0.001) or NS (not significant).

## 3. Discussion

The fish gut microbiome is essential to systemic function, fish health^37^, immune support^38^ and digestive capacity^39^. However, the factors that govern the microbial community colonisation, assembly, and succession in fish species are poorly understood, in terms of both the ecological assembly mechanisms and development within the life cycle. Indeed, in our previous work^23^, we noted a shift in the hindgut microbial community dynamics of juvenile Atlantic cod over time. This displacement was summarised in the current manuscript as a shift from Proteobacteria (mostly *Photobacterium* spp.) to Bacteroidetes and Firmicutes species. Interestingly, this pattern was not observed in fish that found one of the experimental diets unpalatable (ASCO_LG). In this study, we endeavoured to determine the ecological processes that may shed light on this temporal convergence in the hindgut microbiome. We implemented a suite of ‘null-modelling’ approaches to determine the influence of stochastic, deterministic, niche, neutral forces, quantify the contribution of specific ecological processes (selection or dispersal), colonisation, and succession processes. Using the taxa-function robustness measure we determined the potential stability of the gut microbiome to taxonomic perturbation. Understanding such mechanisms is an important step towards managing fish gut microbiome response to newly introduced aquafeed ingredients and improving welfare.

The juvenile cod gut microbiome was driven by deterministic forces. Deterministic forces are those which are shaped by environmental conditions (e.g., pH, nutrient availability), species interactions (e.g., competition and mutualism), or species traits (species genetics). Interestingly, analysis according to Jaccard index (presence/absence) data indicated stochastic processes were driving the microbial community diversity. Thus, we highlight the utility of using the approach by Modin et al. (2020)^40^ to determine the beta-diversity equivalent of hill numbers (^*q*^*d*) by considering the weighting of OTUs in the community. Contrary to our hypothesis, we observed that there was a temporal trend from niche to neutral processes. Neutral processes indicate a lack of environmental selection. This is consistent with Hayes et al. (2020)^41^, where the authors showed neutral processes dominated the gut microbiome in farmed Atlantic salmon (*Salmo salar*). In contrast, however, other authors have found a decreasing influence of neutral processes over time in zebrafish (*Danio rerio*) gut microbial communities^42,43^. However, environmental conditions (e.g. temperature and salinity) are additional variables in these comparisons. That we found multiple assembly processes contributing to overall assembly is unsurprising, since multiple processes can be occurring in a system given the complexity of the biology and interactions with the environment^44^. Moreover, the host environment itself places a deterministic pressure on the microbial communities, given that primarily only species capable of surviving in the gut ecosystem will proliferate in this environment over time^45,46^.

Our results supported evidence of temporal pressure influencing the developing gut microbiome in juvenile cod. In our study, dispersal limitation, variable selection and undominated (weak selection and moderate dispersal) were the primary ecological processes occurring in the cod gut microbiome. The dominant process found varied with respect to time and treatment. For example, variable selection increased at Week 8 for all groups and was the prevailing process at Week 12 in the control group. Variable selection indicates strong selective pressures driving divergent shifts in species composition. However, variable selection was not a dominant process in the macroalgae treatments at Week 12. Yet, the communities in the CTRL, ULVA, and ASCO_N converged in terms of taxonomic composition over the course of the trial. This may point to host-associated factors, however, it is difficult to say with certainty in our study design if patterns are due to host-associated pressures. Host development was found to be a significant influence on driving deterministic microbial community assembly in the zebrafish gut^42^. However, host pressures in the juvenile or adult cod gut are not widely described, a study design similar to the zebrafish study but with Atlantic cod could help elucidate these factors. Research on the Atlantic cod larval phase has suggested that selective pressures in the gut, associated with host intestinal development combined with stochastic pressures, shaped the gut microbiota^47,48^. Ecological theory has been loosely applied to manipulate the microbiome of Atlantic cod larvae by changing the tank water to improve survival rates using selection^49^. How the cod gut microbiota changes through developmental stages is unclear as studies including ours, have focused on a single life stage.

While in the macroalgal dietary treatments dispersal limitation was a core process in the hindgut, this effect was increased in the ASCO_LG fish that found the diet unpalatable. We have not directly cross-compared treatment groups, as each sample represents an individual fish, treatment groups were in separate tanks, and we had sacrificial temporal sampling thus direct dispersal would have been limited between these communities. As defined by Stegen et al. (2015)^50^, dispersal limitation indicates that a low dispersal rate is the primary cause of high compositional turnover (aiding ecological drift or stochastic processes), although the framework does not account for *in situ* diversification. This would occur where dispersal rates are low and new OTUs that evolve within a single taxon may only be present in one community. Given the gut structure whereby the digesta content is contained within the digestive tract (i.e., a semi-closed system), diversification could be an uncharacterised influence here.

The digestive system is a space-limited system, stochastic models such as the competitive lottery model can shed light on what species will be first to occupy the niche space and thus, manipulate the conditions for subsequent species. Interestingly, clades that showed strong lottery-like behaviour (*Alistipes*, Cetobacterium, *Tyzzerella*, and *Fusobacterium*), primarily only did so at Week 0. This indicated that there was a temporal effect and/or that the physical change from a commercial diet to in-house pellets may have impacted microbial competition for space in the hindgut. To fully elucidate this more earlier time points including the former diet would be required (Experiment started Day 366 post hatch baseline prior to changing feed – Week 0). *Photobacterium* species decreased in relative abundance over the course of the feed trial. *Photobacterium* species have a high prevalence in many fish species, including wild cod populations^21,22^. Notably, *Photobacterium* was completely absent of lottery winners (i.e., no OTUs were > 90% within this clade in our samples). Species that do not conform to the competitive lottery schema are thought to be less specialised in their niche and co-exist with other species^31^, which would support the widespread distribution of *Photobacterium* across fish species^19^. Authors have noted that the gut microbiome in Atlantic cod had limited diversity and consisted primarily of *Photobacterium* and other closely related species (showed nepotism)^51^. On this point, the phylogenetic recruitment model showed that the hindgut communities in CTRL and ULVA were neutral while the ASCO_N and ASCO_LG showed overdispersion and under dispersion (more nepotism), respectively. One disadvantage of this study is that we do not have samples for the source metacommunity (tank seawater). In most of the tested community assembly methods, the ASCO_LG fish did not follow the same trend as the other dietary treatments, thus indicating that microbial community assembly patterns were disrupted in fish that displayed poorer growth rates and found the diet unpalatable. These microbial community assembly patterns may drive unexpected changes in fish condition and function^52^. This indicates that diet selection could play a role in the disruption of the microbiome development in farmed fish^53^. Resolving these issues may offer an opportunity to use the gut microbiome to develop fish with improved condition and immune competence for commercial use as in aquaculture or in wild population restocking programmes.

The gut microbiome of any species is integral to the functioning of the system, microbial imbalance can lead to harmful impacts to the host in a condition called dysbiosis^54^. This occurs when the microbial community changes result in a change in function, leading to altered pathways, and the production of excess acids for example. However, the microbial taxonomic composition of a system can fluctuate without detectable changes to the inherent functioning, i.e., functional redundancy, whereby multiple taxa can carry out the same function. We observed taxonomic shifts in the hindgut microbial community over time, and changes in the predicted functional pathways. This may have been related to community imbalance or a natural succession of the communities. In gut microbial communities, the relationship between species taxonomic and functional profiles has been defined as the ‘taxa-function relationship’. This relationship can be viewed as a landscape containing the breadth of microbial taxa and inter-related functional capacities. Using such a landscape, the situations resulting in dysbiosis (microbial imbalance which is harmful to the host) can be assessed. We used the *taxa-function robustness* measure^55^ to calculate the breadth of taxonomic shifts (perturbations) and the community functional capacity. In our work, we showed that in the hindgut microbiome, despite temporal taxonomic shifts, the microbial communities increased in robustness and functional stability, particularly for functions involved in metabolism and cell regulation over time. The community exhibited greater functional redundancy over time, indicating an overlap of functional ability. In general, the ASCO_LG individuals followed a different trend. These findings could be improved with the addition of metagenomic data to provide a less predictive approach. This work highlights that functional analysis of microbial communities in complex systems, such as the gut has a greater utility than taxonomic profiles^56^.

In summary, the ecological drivers of microbial community assembly in the gut microbiome are important factors to consider when linking microbial community composition and diversity to fish health and environmental parameters. We conclude that the microbiome in the gut of Atlantic cod (*G. morhua*) in an experimental feeding trial over a twelve-week period was under the influence of multiple assembly processes (deterministic assembly and a trend from niche to neutral processes). We quantified these processes as an increase in variable selection in the control diet (divergence in communities related to selective environmental conditions) over time. Dispersal limitation was a driver of the gut microbiome for fish fed the macroalgae supplemented diet at Week 12. Clades that conformed to the competitive lottery schema and had ‘winning’ OTUs (*Alistipes*, Cetobacterium, *Tyzzerella*, and *Fusobacterium*) only showed status behaviour at Week 0. *Photobacterium*, an important taxon in fish gut research did not show lottery-like behaviour. The recruitment of new taxa overtime was altered in the ASCO (10% *Ascophyllum nodosum* supplement), with individuals who found the diet unpalatable exhibiting phylogenetic under dispersion (nepotistic recruitment of species). Finally, the gut microbiome showed increasing robustness to taxonomic disturbance over time and an increase in functional redundancy, except for the ASCO_LG individuals. These fish showed an altered microbiome, with increased susceptibility to functions related to infectious diseases and cell regulation. Finally, although our study focused on the juvenile cod gut microbiome, many of these findings are of broad interest in fish research, and indeed to the wider field of gut microbiome research. Further research is needed to unravel the complex interactions between host and microbiome to determine how it influences strong selection on the gut microbiome. Targeted ‘omic sequencing and metabolomics of the digesta content to track the gut microbiome and function over the complete life cycle of Atlantic cod (*G. morhua*) may elucidate these factors. Such models that may emerge could be applied to other important marine fish species for better health management in terms of prebiotics and probiotics as functional feed additives.

## 4. Materials and Methods

A full description of the experimental design, fish feed formulation, fish condition, sample collection and sequencing analysis is included in our previous work^23^. Briefly, juvenile Atlantic cod (*G. morhua*) were hand-graded (123 ± 7 g, SD) and randomly allocated to one of nine experimental tanks (60 individuals per tank). Fish were acclimated for one week on a commercial fishmeal diet (Amber Neptun, Skretting, Stavanger, Norway), noted as the Week 0 sampling phase. Then tanks were assigned to the following diets, a 10% dietary macroalgae supplement (either *Ulva rigida* [ULVA] or *Ascophyllum nodosum* [ASCO] species) or a control diet for basal comparison, i.e., no algal addition [CTRL]. At Week 8 within the feed trial, a subgroup of the ASCO fish displaying ‘reduced growth rates were observed and was likely due to reduced acceptance of the feed^23^. We therefore subcategorised this group as [ASCO_LG], and the remaining fish with ‘normal’ growth were referred to as [ASCO_N] (see Keating et al 2021^23^ for further discussion). The feed trial was carried out for twelve weeks.

### 4.1 Sample Collection, DNA extraction and 16S rRNA amplicon sequencing

At each noted time-point: Week 0, Week 8, and Week 12, individual fish were removed from the tanks and euthanised with an overdose of tricaine methanesulphonate solution (MS222, Pharmaq, Overhalla, Norway). The brain was then destroyed to confirm death according to regulations for animal welfare (EU Directive 2010/63/EU). The digestive tract of the fish was aseptically removed and the digesta content from the last 10-15% of the digestive tract was aliquoted into sterile microcentrifuge tubes. Samples were transported to the National University of Ireland Galway on dry ice and stored at -80□C. DNA extractions from hindgut digesta per sample were carried out using a modified phenol-chloroform extraction method that included a bead-beating step using Lysing Matrix E tubes (MP Biomedical, Illkirch-Graffenstaden, France), as described previously^23,57^. Sample DNA and DNA from a negative extraction control (nuclease-free water, Qiagen, Venlo, The Netherlands) were sent to the Research Technology Support Facility at Michigan State University (Michigan, USA) for sequencing. Amplicon sequencing of the 16S rRNA gene targeting the V4 hypervariable region using the universal primer set [515f/806r^58^]. Sequencing was carried out using the Illumina technology using a standard flow cell and 500 cycle v2 reagent cartridge (Illumina Inc., Hayward, California, USA).

### 4.2 Bioinformatics

The methods to generate all microbial data (e.g., operational taxonomic units (OTUs), phylogenetic trees and biom. files) are given in Keating et al (2021)^23^. The raw sequences are also available in the SRA database under Bioproject Submission PRJNA636649.

### 4.3 Subset Analysis

For finding key microbial species that contribute to beta-diversity between samples over time we used the “BVSTEP” routine^59^. Briefly, the method calculates the Bray-Curtis distance between samples using all the OTUs and records it as original distances. It then permutes through the subset of OTUs, and for each permutation, it calculates the Bray-Curtis distances between the samples again and correlates these distances against the original recorded distances until subsets are obtained that explain roughly the same beta diversity as the full set of OTUs. To run this algorithm, bvStep() (from the sinkr package) was used^60^. After obtaining the subset of OTUs, we used R’s ‘Vegan’ package^61^, particularly the bioenv () function to regress the subsets on top of the principal coordinate analysis (PCOA) plot.

### 4.4 Microbial Community Assembly

#### 4.4.1. Hill-numbers dissimilarity indices (^*q*^*d*)

Hill numbers are a set of indices parameterised by q representing the diversity order which determines the weight given to the relative abundance of OTUs in a community. ^40^ derived the beta-diversity equivalent of hill numbers (^*q*^*d*) as a dissimilarity index of diversity order where ^*q*^*d* values are scaled between 0 (similar) and 1 (dissimilar). The authors used this approach to illustrate how OTU abundance contributes to the dissimilarity between communities. Here, we compared ^*q*^*d* at q=0 (presence/absence, i.e., Jaccard index), q = 1 (OTUs abundances are weighted ie Bray-Curtis) and q = >1 (OTUs with greater relative abundance have increased weighting). Further, a randomisation scheme was applied, and repeated many times to obtain the null distribution for the dissimilarities across this scale between communities. These null distributions when compared to the observed dissimilarity (^*q*^*d*) reveal ecological insights, i.e., if the values are similar, the observed dissimilarity can be explained by stochastic factors, and if higher or lower than the null expectation, then there are likely deterministic factors that favour different or similar microbial taxa in two categories. These indices and null distributions were calculated using the qdiv Python software^40^.

#### 4.4.2 Beta-Null Model

Beta-null deviation measures were calculated according to Tucker et al (2016) and Lee et al. (2017)^35,62^. The method first calculates the pairwise observed dissimilarities □_obs_) between samples using the Generalised Unifrac dissimilarity measure. By preserving the alpha diversity in the observed samples, random communities are generated (999 randomisations) to calculate the beta diversity measure again for these communities □_null_) and then the deviation from the observed dissimilarities are recorded. The average of the deviations gives a numerical value that differentiates between niche (values further from 0) and neutrally structured communities (values nearer to 0). We have done this separately for each treatment group on a temporal basis (Week 0, CTRL_08, ULVA_08, ASCO_N_08, ASCO_LG_08, CTRL_12, ULVA_12, ASCO_N_12, and ASCO_LG_12).

#### 4.4.3 Quantitative Process Elements

Quantitative process elements (QPE) were used to assess the influence of ecological processes based on the conceptual framework of Vellend et al (2010)^63^ and implemented according to Stegen et al (2013, 2015)^50,64^. This framework provides a quantitative measure of the influence of selection and dispersal pressures on microbial community structure. Selection considers deterministic selective pressure which results in divergent (variable selection) or convergent (homogenous selection) communities often considered overtime. While dispersal considers the spatial movement of species where the increased movement of species results in convergent communities (homogenising dispersal), or limited movement of species results in divergent communities through drift (dispersal limitation). The authors also included the category ‘Undominated’ to describe the situation whereby neither selection nor dispersal processes dominate.

The framework considers the phylogenetic distance and phylogenetic turnover between closely related OTUs in pairwise samples^29^. This is achieved using the abundance-weighted β-mean-nearest taxon distance (βMNTD)^65^. To determine how this varied from the null expectation, randomisations were employed whereby the abundances and species names were shuffled across phylogenies to provide a null value^64^. This was replicated 999 times to give the null distribution. The deviation between the null distribution and the observed βMNTD value = β-nearest taxon index (βNTI). If the observed βMNTD value is significantly greater (βNTI > 2) or less (βNTI < −2) than the null expectation, the community is assembled by variable or homogeneous selection, respectively. For the remaining with no significant deviation, in the next step, Raup-Crick was used with the inclusion of OTU relative abundance ^66^ termed RC_bray_. RC_bray_ values were compared to the null expectation and the resulting deviation determined the influence of dispersal (RC_bray_ > .95). Values of RC_bray_ > -.95 indicate *Homogenising Dispersal* (transport between microbiomes leading to establishment), while values of RC_bray_ > +.95 indicate *Dispersal Limitation*. In the latter, this may indicate ‘true’ effects of dispersal limitation (i.e., limited transport across microbiomes and stochastic events) and/or historical contingency. In cases where values were <0.95 the communities were ‘Undominated’, i.e., not dominated by a sole ecological process.

#### 4.4.4. Competitive Lottery Model

We applied the competitive lottery model as outlined in^31^ which is based on the theory of Sale (1977)^67^ first proposed for fish populations. The theory is based on the idea that there is competition between colonising species within a niche and only a single species can ‘win’ in the space (strong priority effect). This ‘winner’ is chosen at random (stochastic process) with an analogy drawn to a ‘lottery’. In microbial ecology this scheme determines clade-based assembly, i.e., within a taxonomic group (a genus), we can determine if the group follows lottery-like behaviour and if so what OTUs ‘win’. A winning species/OTU is defined as a clade member with >90% abundance [see Verster and Borenstein (2018)^31^] for details on how this threshold was determined. Then the diversity of lottery winners was calculated using Shannon diversity index of the winners across samples (how often each OTU occurs as a winner in samples where lottery-like behaviour was observed). Diversity was normalised between 0 and 1 to account for differences in lottery winners. Values approaching 0 indicate that a single OTU is dominating that specific genus in all samples, while values approaching 1 indicate an even distribution of winning OTUs within a genus.

#### 4.4.5. Phylogenetic Recruitment Model

The phylogenetic recruitment model^68^ was then used to describe the order in which new species are detected in the cod hindgut microbiome over time. In this model, the dispersion parameter (D) is calculated based on the probability of detection of new species on temporal scales by fitting a logistic error model on changes in phylogenetic diversity (PD) estimates. As opposed to previous models, time-series dependency is assumed. The value of D determines the community assembly mechanisms with D = 0 indicating the neutral model (all species have an equiprobable chance of detection). The community is said to be over dispersed when D > 0. In this case, species from the species pool have higher probabilities of detection when they are more phylogenetically divergent from the species that have already been detected. In contrast, the community is said to be under dispersed when D < 0. Here, species from the species pool have higher probabilities of detection when they are more phylogenetically similar to species that have already been detected (exhibiting nepotism).

### 4.5. Functional Analysis and Taxa-Function Robustness

#### 4.5.1 Functional Analysis

The functional potential was obtained as KEGG orthologs (KO) and pathway predictions by using Phylogenetic Investigation of Communities by Reconstruction of Unobserved States (PICRUSt2)^69^ and the Quantitative Insights into Microbial Ecology (QIIME2) plugin^70^ using the parameters --p-hsp-method pic --p-max-nsti 2. To find KEGG enzymes/MetaCyc pathways that are significantly different between different categories, we used DESeqDataSetFromMatrix() function from DESeq2^71^ package with the adjusted p-value significance cut-off of 0.05 and log_2_ fold change cut-off of 2. This function uses a negative binomial general linear model to obtain maximum likelihood estimates for OTUs log fold change between two conditions. Then Bayesian shrinkage is applied to obtain shrunken log-fold changes subsequently employing the Wald test for obtaining significances. For KEGG orthologs that were at least log_2_ fold significant, we used iPath3^72^ to give an overview of KEGG pathways for microbial metabolic function.

#### 4.5.2. Taxa-Function Robustness

Following the procedure of Eng and Borenstein (2018)^55^, the *taxa-function robustness* measure of the cod hindgut microbial communities was calculated. The principle of the taxa-function robustness measure is to perturb an individual sample several times (100 perturbations) and then calculate a two-dimensional profile of taxonomic shift versus functional shift. To create the taxonomic profiles, weighted Unifrac dissimilarities were calculated across samples and simulated perturbations. To obtain predicted functional profiles, the authors calculated the KEGG Orthology (KO) groups for the whole green genes database (gg_13_5) along with KO copy numbers provided as a reference database (https://github.com/borenstein-lab/robustness), which can be used if the OTUs follow the green genes nomenclature. For the functional profiles, cosine dissimilarities were calculated across samples and simulated perturbations. After obtaining the taxonomic and functional shifts (both denoted as *t*) for a given sample, a relationship between taxonomic perturbation magnitude and functional profile shift is assumed to behave individually as 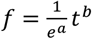 and fitted using the linear regression model on natural log transformed data: In(*f*) = −*a* + *b* In(*t*). In the equation, *f* denotes the expected shift in functional profile and we can estimate **a** (termed as “attenuation” coefficient describing *the expected rate at which increases in the taxonomic perturbation magnitude are expected to increase functional profile shifts*) and **b** (termed as “buffering” coefficient indicating how large a perturbation must be before a functional profile shift becomes noticeable). The coefficients thus serve as proxies (robustness factors) to summarise the property of a sample to withstand perturbation. These were then calculated for all the samples in the dataset. Additionally, the main gene distribution features (GDFs) across the genomes of species in a community, were then displayed as a PCOA plot. Further details can be found in Eng and Borenstein (2018)^55^.

## 5. Data Availability Statement

The raw sequences are also available in the SRA database under Bioproject Submission PRJNA636649.

## Acknowledgements

We dedicate this paper to Dr Richard Fitzgerald, a dear colleague and friend, 1957-2016. The authors also kindly acknowledge all staff at the Carna Research Station, National University of Ireland Galway, Carna, Co. Galway, Ireland for their technical assistance during the feed trial.

## 9. Author Contributions

CK dissected the fish for microbiome sampling, DNA extraction optimisation and extractions on the gut content, bioinformatics, statistical analysis and data interpretation. MBW managed the feeding trial, coordinated sampling and data management of tank populations. JH and RD ran the feeding trial and assisted with sampling. SW ran the feeding trial and assisted with dissections for microbiome analysis. AHLW was responsible for seaweed identification and collection, designed the test diets and feed analysis. UZI wrote the analysis scripts and performed the microbiota bioinformatic and statistical analysis, and data interpretation with CK. UZI and CK conceived the ecological assembly research question study. While CJS, RF, MBW, SJD and AHLW designed the experimental feeding-trial study. CK and UZI wrote the manuscript, which was reviewed and edited by all authors. RF was EIRCOD Project Coordinator. All authors made a substantial contribution towards the research study and all authors (except RF) approved the final manuscript.

## 12. Supplementary Information

**Figure S1.**
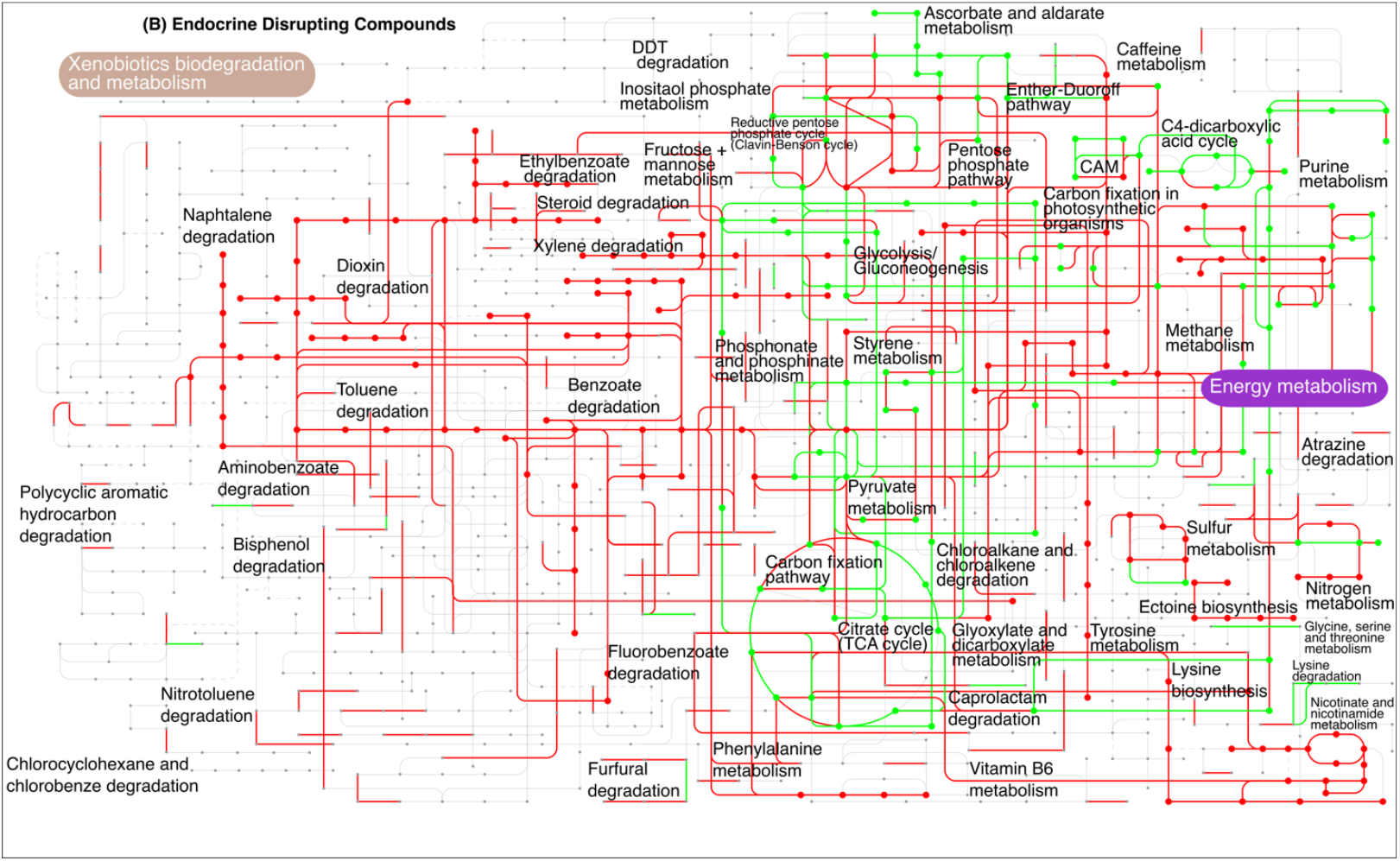
KEGG orthologs and pathway predictions from the 16S rRNA gene sequencing of the gut microbiome from juvenile Atlantic cod using PICRUSt2 predictive functions^69^. The figure shows the Metacyc pathways drawn in iPath3^72^ which are significantly different at a log_2_ fold change (P < 0.05) from Week 0 (red) and Week 12 (green).

**Figure S2.**
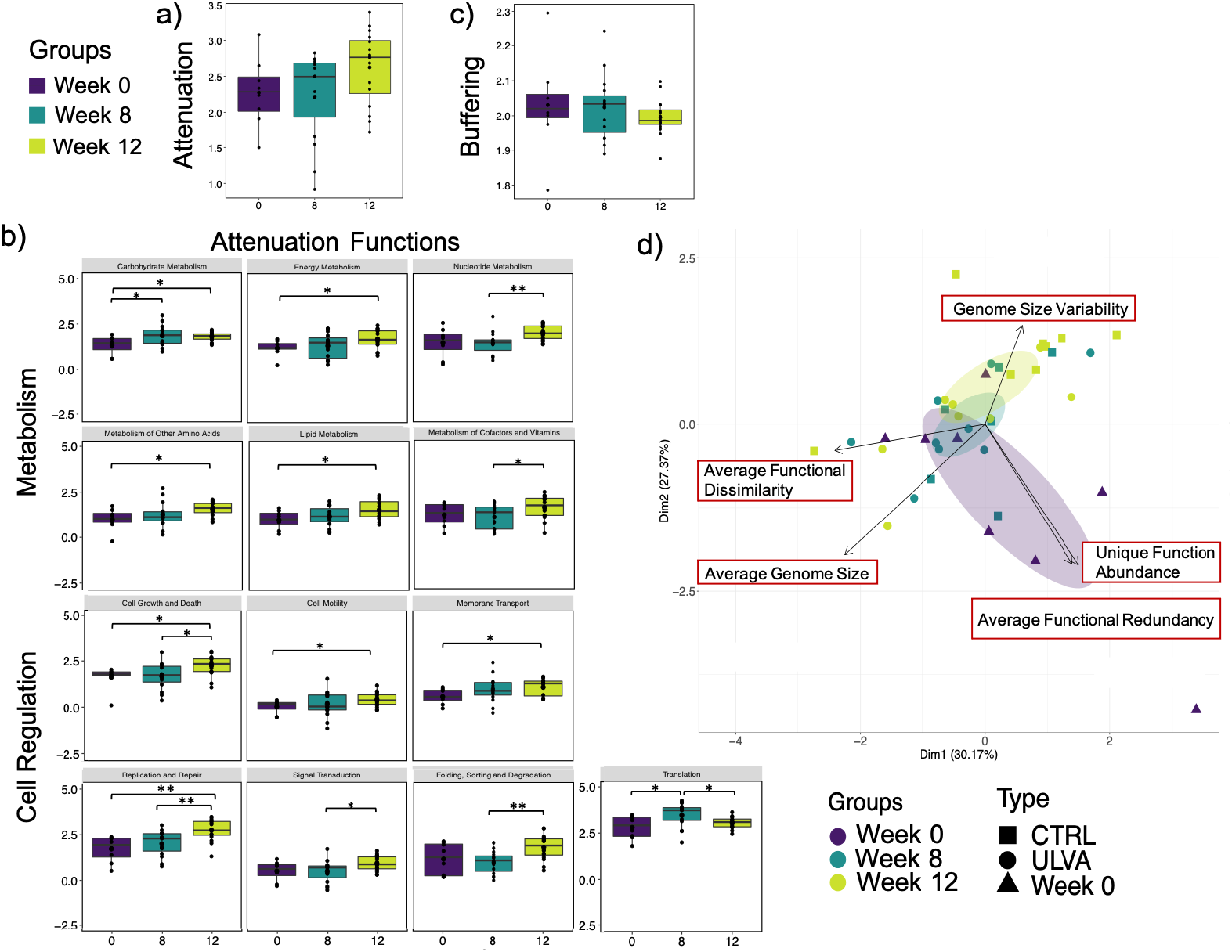
Taxa-function robustness in the Atlantic cod hindgut microbiome compared across time excluding the ASCO samples (Groups: Week 0, Week 8 [CTRL and ULVA] and Week 12 [CTRL and ULVA]). **a)** The attenuation values for each time group. **b)** The specific-robustness attenuation functions that were statistically significant between time groups. **c)** The buffering values for each time group. **d)** Principal co-ordinate analysis plot of the five gene distribution factors compared across time group (Week 0, Week 8 and Week 12). Treatment group is indicated in symbols (Square = CTRL, Circle = ULVA, and Triangle (Week 0). Lines connecting categories shows were significant (ANOVA) with * (p < 0.05), ** (p < 0.01), or *** (p < 0.001).

**Table S1.** Table showing the ‘lottery’ winners in genus clade groups (OTUs with > 90% within the clade). The table shows the genus name, time group, winner prevalence in samples and the normalised winner diversity and the identity of the winning OTUs.

| Clade | Groups | Winner prevalence | Normalised Winner diversity | Winners |
| --- | --- | --- | --- | --- |
| *Rikenella* | 0 | 0.5 | 0 | OTU_41: Bacteria; Bacteroidetes; Bacteroidia; Bacteroidales; Rikenellaceae; Rikenella; |
| *Rikenella* | 8 | 0.666666667 | 0 | OTU_41: Bacteria; Bacteroidetes; Bacteroidia; Bacteroidales; Rikenellaceae; Rikenella; |
| *Rikenella* | 12 | 0.625 | 0.295903274 | OTU_41: Bacteria; Bacteroidetes; Bacteroidia; Bacteroidales; Rikenellaceae; Rikenella;<br>OTU_37: Bacteria; Bacteroidetes; Bacteroidia; Bacteroidales; Rikenellaceae; Rikenella; |
| *Bacteroides* | 0 | 0.857142857 | 0.231542658 | OTU_9: Bacteria; Bacteroidetes; Bacteroidia; Bacteroidales; Bacteroidaceae; Bacteroides;<br>OTU_2969: Bacteria; Bacteroidetes; Bacteroidia; Bacteroidales; Bacteroidaceae; Bacteroides; |
| *Bacteroides* | 8 | 0.666666667 | 0 | OTU_9: Bacteria; Bacteroidetes; Bacteroidia; Bacteroidales; Bacteroidaceae; Bacteroides; |
| *Bacteroides* | 12 | 0.466666667 | 0 | OTU_9: Bacteria; Bacteroidetes; Bacteroidia; Bacteroidales; Bacteroidaceae; Bacteroides; |
| *Tyzzarella* | 0 | 0.833333333 | 0.360964047 | OTU_23: Bacteria; Firmicutes; Clostridia; Clostridiales; Lachnospiraceae; Tyzzarella;<br>OTU_153: Bacteria; Firmicutes; Clostridia; Clostridiales; Lachnospiraceae; Tyzzarella; |
| *Tyzzarella* | 8 | 0.8 | 0.271782222 | OTU_23: Bacteria; Firmicutes; Clostridia; Clostridiales; Lachnospiraceae; Tyzzarella;<br>OTU_153: Bacteria; Firmicutes; Clostridia; Clostridiales; Lachnospiraceae; Tyzzarella; |
| *Tyzzarella* | 12 | 0.714285714 | 0.234497797 | OTU_23: Bacteria; Firmicutes; Clostridia; Clostridiales; Lachnospiraceae; Tyzzarella;<br>OTU_90: Bacteria; Firmicutes; Clostridia; Clostridiales; Lachnospiraceae; Tyzzarella; |
| *Macellibacteroides* | 0 | 0.5 | 0 | OTU_2550: Bacteria; Bacteroidetes; Bacteroidia; Bacteroidales; Porphyromonadaceae; Macellibacteroides; |
| *Macellibacteroides* | 8 | 0.25 | 0 | OTU_2550: Bacteria; Bacteroidetes; Bacteroidia; Bacteroidales; Porphyromonadaceae; Macellibacteroides; |
| *Macellibacteroides* | 12 | 0.066666667 | 0 | OTU_2550: Bacteria; Bacteroidetes; Bacteroidia; Bacteroidales; Porphyromonadaceae; Macellibacteroides; |
| *Fusobacterium* | 0 | 0.833333333 | 0.455485915 | OTU_3155: Bacteria; Fusobacteria; Fusobacteriia; Fusobacteriales; Fusobacteriaceae; Fusobacterium;<br>OTU_2525: Bacteria; Fusobacteria; Fusobacteriia; Fusobacteriales; Fusobacteriaceae; Fusobacterium; |
| *Fusobacterium* | 8 | 0.416666667 | 0.455485915 | OTU_2525: Bacteria; Fusobacteria; Fusobacteriia; Fusobacteriales; Fusobacteriaceae; Fusobacterium; |
|  |  |  |  | OTU_3155: Bacteria; Fusobacteria; Fusobacteriia; Fusobacteriales; Fusobacteriaceae; Fusobacterium; |
| *Fusobacterium* | 12 | 0.142857143 | 0 | OTU_2525: Bacteria; Fusobacteria; Fusobacteriia; Fusobacteriales; Fusobacteriaceae; Fusobacterium; |
| *Lachnoclostridium* | 0 | 0.8 | 0 | OTU_2624: Bacteria; Firmicutes; Clostridia; Clostridiales; Lachnospiraceae; Lachnoclostridium; |
| *Lachnoclostridium* | 8 | 0.285714286 | 0.5 | OTU_2624: Bacteria; Firmicutes; Clostridia; Clostridiales; Lachnospiraceae; Lachnoclostridium;<br>OTU_130: Bacteria; Firmicutes; Clostridia; Clostridiales; Lachnospiraceae; Lachnoclostridium; |
| *Lachnoclostridium* | 12 | 0.5 | 0.271782222 | OTU_2624: Bacteria; Firmicutes; Clostridia; Clostridiales; Lachnospiraceae; Lachnoclostridium;<br>OTU_130: Bacteria; Firmicutes; Clostridia; Clostridiales; Lachnospiraceae; Lachnoclostridium; |
| *Alistipes* | 0 | 0.8 | 0.646014837 | OTU_34: Bacteria; Bacteroidetes; Bacteroidia; Bacteroidales; Rikenellaceae; Alistipes;<br>OTU_20: Bacteria; Bacteroidetes; Bacteroidia; Bacteroidales; Rikenellaceae; Alistipes;<br>OTU_3429: Bacteria; Bacteroidetes; Bacteroidia; Bacteroidales; Rikenellaceae; Alistipes; |
| *Alistipes* | 8 | 0.25 | 0 | OTU_3429: Bacteria; Bacteroidetes; Bacteroidia; Bacteroidales; Rikenellaceae; Alistipes; |
| *Alistipes* | 12 | 0.066666667 | 0 | OTU_62: Bacteria; Bacteroidetes; Bacteroidia; Bacteroidales; Rikenellaceae; Alistipes; |
| *Erysipelatoclostridium* | 8 | 0.166666667 | 0 | OTU_2180: Bacteria; Firmicutes; Erysipelotrichia; Erysipelotrichales; Erysipelotrichaceae; Erysipelatoclostridium; |
| *Cetobacterium* | 0 | 1 | 0.459147917 | OTU_2113: Bacteria; Fusobacteria; Fusobacteriia; Fusobacteriales; Fusobacteriaceae; Cetobacterium;<br>OTU_355: Bacteria; Fusobacteria; Fusobacteriia; Fusobacteriales; Fusobacteriaceae; Cetobacterium; |
| *Cetobacterium* | 8 | 0.5 | 0 | OTU_2113: Bacteria; Fusobacteria; Fusobacteriia; Fusobacteriales; Fusobacteriaceae; Cetobacterium; |
| *Cetobacterium* | 12 | 0.444444444 | 0.75 | OTU_3117: Bacteria; Fusobacteria; Fusobacteriia; Fusobacteriales; Fusobacteriaceae; Cetobacterium;<br>OTU_355: Bacteria; Fusobacteria; Fusobacteriia; Fusobacteriales; Fusobacteriaceae; Cetobacterium;<br>OTU_2113: Bacteria; Fusobacteria; Fusobacteriia; Fusobacteriales; Fusobacteriaceae; Cetobacterium; |
| *[Anaerorhabdus] furcosa group* | 0 | 0.285714286 | 0 | OTU_3118: Bacteria; Firmicutes; Erysipelotrichia; Erysipelotrichales; Erysipelotrichaceae; [Anaerorhabdus] furcosa group; Erysipelotrichaceae feline oral taxon 121; |
| *[Anaerorhabdus] furcosa group* | 8 | 0.222222222 | 0 | OTU_3118: Bacteria; Firmicutes; Erysipelotrichia; Erysipelotrichales; Erysipelotrichaceae; [Anaerorhabdus] furcosa group; Erysipelotrichaceae feline oral taxon 121; |
| *[Anaerorhabdus]*<br>*furcosa group* | 12 | 0.36363636<br>4 | 0.51185950<br>7 | OTU_3282: Bacteria; Firmicutes; Erysipelotrichia; Erysipelotrichales; Erysipelotrichaceae; [Anaerorhabdus] furcosa group;<br>OTU_3118: Bacteria; Firmicutes; Erysipelotrichia; Erysipelotrichales; Erysipelotrichaceae; [Anaerorhabdus] furcosa group; Erysipelotrichaceae feline oral taxon 121; |

